# Spinal stretch reflexes exploit musculoskeletal redundancy to support postural hand control

**DOI:** 10.1101/270116

**Authors:** Jeffrey Weiler, Paul L Gribble, J. Andrew Pruszynski

## Abstract

Motor behaviour is most efficiently controlled by only correcting disturbances or deviations that influence task success. It is currently thought that such sophisticated control is computed within a transcortical feedback pathway. Here we show that even the fastest spinal feedback pathway can produce corrective responses that adhere to this control scheme. We first applied small mechanical perturbations that flexed the elbow joint – stretching the triceps muscle – and simultaneously flexed or extended the wrist joint, displacing the hand various distances away from a central target. We then changed the arm’s orientation and applied the same joint perturbations, which reversed the mapping between joint motion and hand displacement. In all cases, we found that the triceps’ spinal stretch reflex was tuned to the hand’s displacement relative to the target, and not how the triceps muscle was stretched. Our findings reveal that the fastest spinal feedback pathway is capable of integrating and modulating feedback from multiple muscles to produce efficient corrective responses, forcing a re-evaluation of the how the nervous system derives the sophisticated control laws that support natural motor behaviour.

## Introduction

Real-world actions require actively controlling many joints in the presence of internal and external disturbances (Faisal et al, 2008). The simplest way for the nervous system to counteract disturbances is to ensure that all the joints remain at some specific set of reference positions by independently correcting deviations at each joint. However, a better way for the nervous system to counteract disturbances is by taking advantage of musculoskeletal redundancy and adhering to the so-called minimum intervention principle – that is, correcting only joint deviations to the degree that they interfere with task success (Todorov 2004; Todorov and Jordan, 2002).

Many behavioural studies have shown that the nervous system adheres to the minimum intervention principle (Cole and Abbs, 1987; Diedrichsen, 2007; Dimitriou et al, 2012; Gracco and Abbs, 1985; Mutha and Sainburg, 2009; Omrani et al, 2013; Robertson and Miall, 1997; Scholz et al, 2000). For example, when people reach to grasp an object, errors introduced by experimentally perturbing one finger are not only corrected by responses at the perturbed finger but at all the fingers that help govern grasp aperture (Cole, Gracco and Abbs, 1984). A key outstanding question in sensorimotor neuroscience is which neural circuits implement the sophisticated control laws that produce such behaviour (Scott 2004; Scott 2016). Sixty years of work, primarily focusing on reaching actions, indicates that spinal circuits may not perform the requisite computations and that this capacity may be a specialization of a transcortical feedback pathway through primary motor cortex and other cortical regions involved in the production of voluntary movement (Cheney and Fetz 1984; Evarts and Fromm 1977; Evarts and Tanji 1976; Omrani et al, 2014, 2016; Picard and Smith 1992; Pruszynski et al, 2011, 2012, 2014; Wolpaw 1980).

Here we show that, in the context of postural hand control, even the fastest spinal feedback pathway can produce solutions consistent with the minimum intervention principle. In our first experiment, participants maintained their hand at a spatial target while we applied small mechanical perturbations to their elbow and wrist joints. We chose mechanical perturbations that moved the participant’s hand away from the target to varying degrees, but critically, we ensured that the perturbation that yielded the largest hand displacement did so with the least elbow rotation. Consistent with the minimum intervention principle, we found that spinal stretch reflexes at the elbow were tuned to hand displacement relative to the target, rather than the amount of elbow rotation. In our second experiment, we dissociated wrist rotation from how the hand moved relative to the target by having participants adopt two different arm orientations. In this arrangement, the same mechanical perturbation at the wrist moved the hand away from the target in one arm orientation but towards the target in the other arm orientation. We again found that spinal stretch reflexes at the elbow were tuned to hand displacement rather than elbow rotation. In fact, changing the arm’s orientation completely reversed the pattern of spinal stretch reflexes at the elbow in a way that was appropriate for returning the hand to the target. Taken together, these findings reveal that this spinal feedback pathway is more sophisticated than previously thought – capable of integrating and modulating feedback from multiple muscles to produce corrective responses that take advantage of musculoskeletal redundancy.

## Results

### The spinal stretch reflex accounts for hand displacement

In our first experiment, participants (n = 25) grasped the handle of a robotic exoskeleton and placed their hand at a central target. The robot then mechanically flexed their elbow, stretching the triceps muscle, and simultaneously flexed, extended, or did not alter the angle of their wrist (**Fig. 1A**). All of the mechanical perturbations moved the participant’s hand away from the target. Critically, how far that hand was displaced from the target was a function of both the wrist and elbow perturbation (**Fig. 1B-D**).

**Figure 1.**
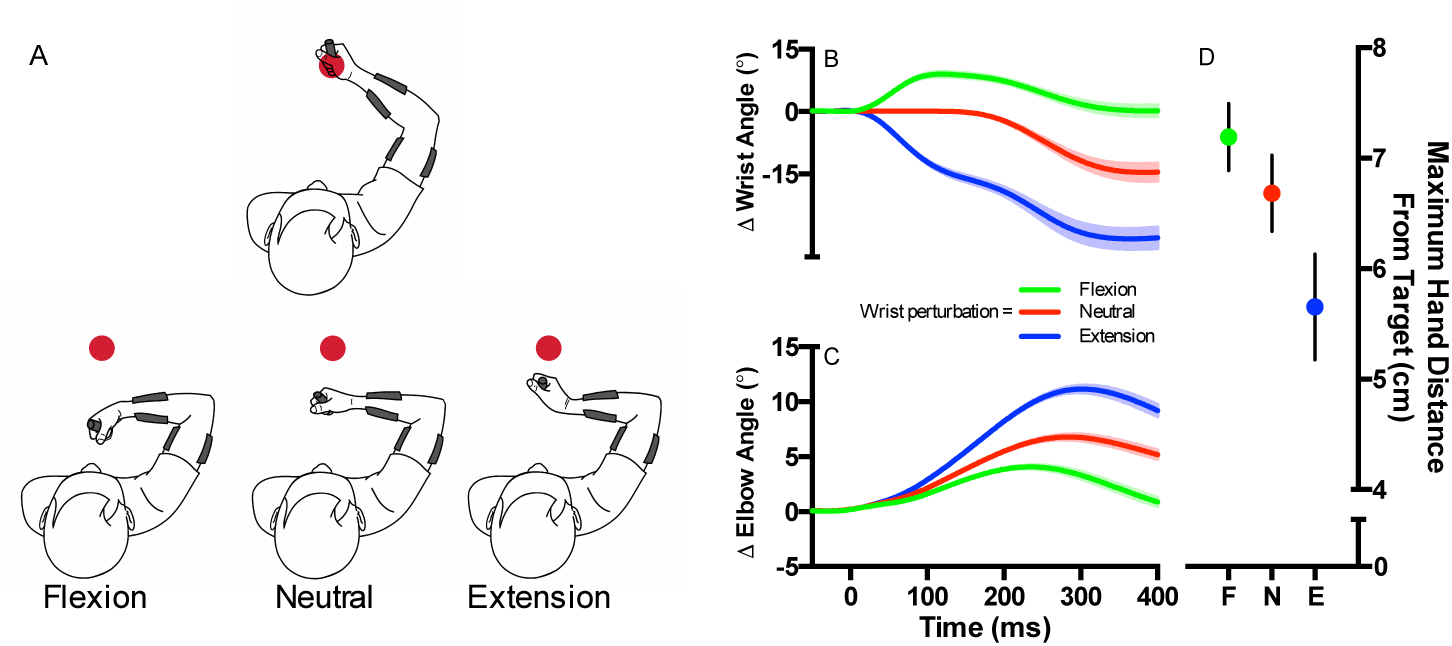
A: Cartoon showing how the elbow flexion perturbation and simultaneous wrist flexion perturbation (left), no wrist perturbation (center) and wrist extension perturbation (right) displaced the hand from the target (red dot). B: Mean change in wrist angle aligned to perturbation onset. Green, and blue traces reflect perturbations that flexed and extended the wrist, respectively. Red trace reflects trials in which no mechanical perturbation was applied to the wrist. Shading reflects ±1 SEM. C: Same format as B, but for elbow angle. D: Maximum hand displacement from the target. The dots represent the group mean for the three wrist perturbation conditions (F = flexion; N = none; E = extension). Error bars reflect ±1 SEM.

Participants were instructed to counteract the perturbation and return their hand to the target quickly and accurately. They did so not by simply returning their joints to their initial positions, but by simultaneously extending their elbow and wrist joints in a coordinated fashion while keeping their shoulder joint relatively fixed – a behaviour that is consistent with the minimum intervention principle (**Fig. 1B**,**C, for shoulder kinematics see Sup Fig. 1**). We found that the triceps’ spinal stretch reflex (i.e., mean EMG activity 25-50ms post perturbation) was tuned to the distance the hand was displaced from the target, and not the amount of elbow flexion, (F(2,28) = 103.5, p < 0.001; post-hoc trend analysis: linear F(1,24) = 127.04, p < 0.0001; quadratic F(1,24) = 2.05, p = 0.17). In fact, the magnitude of the triceps’ spinal stretch reflex was largest when the triceps muscle was stretched the least and even inhibited relative to baseline when it was stretched the most (**Fig. 2**).

**Figure 2.**
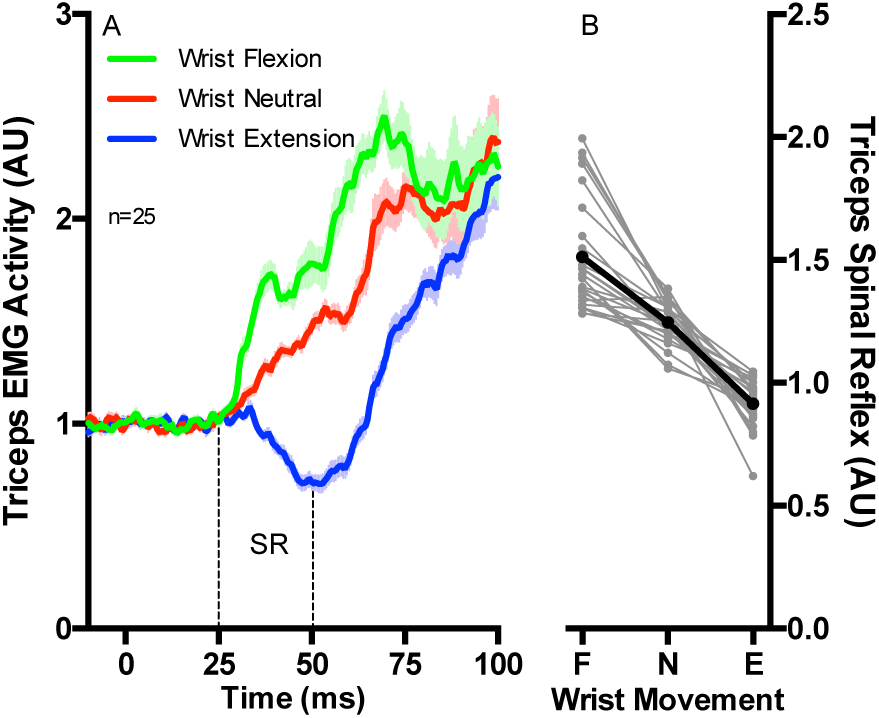
A: Mean triceps EMG activity following the mechanical perturbation. Green and blue traces reflect wrist perturbations that flexed and extended the wrist, respectively, whereas the red trace reflects trials in which no perturbation was applied to the wrist. Data is aligned to perturbation onset. Shading reflects ±1 SEM. The spinal stretch reflex epoch (SR) spans from 25-50 ms relative to perturbation onset. B: Mean triceps EMG activity in the spinal stretch reflex epoch for the three wrist perturbations (F = flexion; N = none; E = extension). Thin grey lines reflect individual participants and the thick black lines reflect the group mean.

### Volitional intent does not modify the tuning of the spinal stretch reflex

In the above analysis we showed that the triceps’ spinal stretch reflex was modulated based on the magnitude of elbow extension needed to return the hand to the target, rather than the amount the triceps was stretched. These results run counter to a long history of experiments showing that spinal stretch reflexes are not modulated by an individual’s volitional intent. To test the role of volitional intent we instructed a sub-set (n = 15) of the participants from our first experiment to complete an additional block of trials in which they were told to ‘not intervene’ following the mechanical perturbation.

What is typically observed in the context of this manipulation is that spinal stretch reflexes are not modulated by instruction, whereas responses that include inputs from the transcortical feedback pathway (i.e., the long-latency stretch reflex: muscle activity 50-100 ms post perturbation) are influenced by instruction (see Hammond, 1956). Our data are consistent with this classical finding. Specifically, the magnitude of the triceps’ spinal stretch reflex was not influenced by instruction, neither when the wrist was flexed, not perturbed, nor extended (ts(14) all < 1.58, ps all > 0.135). In contrast, the triceps’ long-latency stretch reflex was influenced by instruction, and this occurred for all three wrist perturbation conditions (ts(14) all > 4.01, ps all < 0.001: repeated measures ANOVA three-way interaction [epoch (spinal, long-latency) by wrist perturbation (flexed, neutral, extended) by volitional intent (counteract, do not intervene)] for initial omnibus test (F(2,28) = 17.19, p < 0.001) (**Fig. 3**).

**Figure 3.**
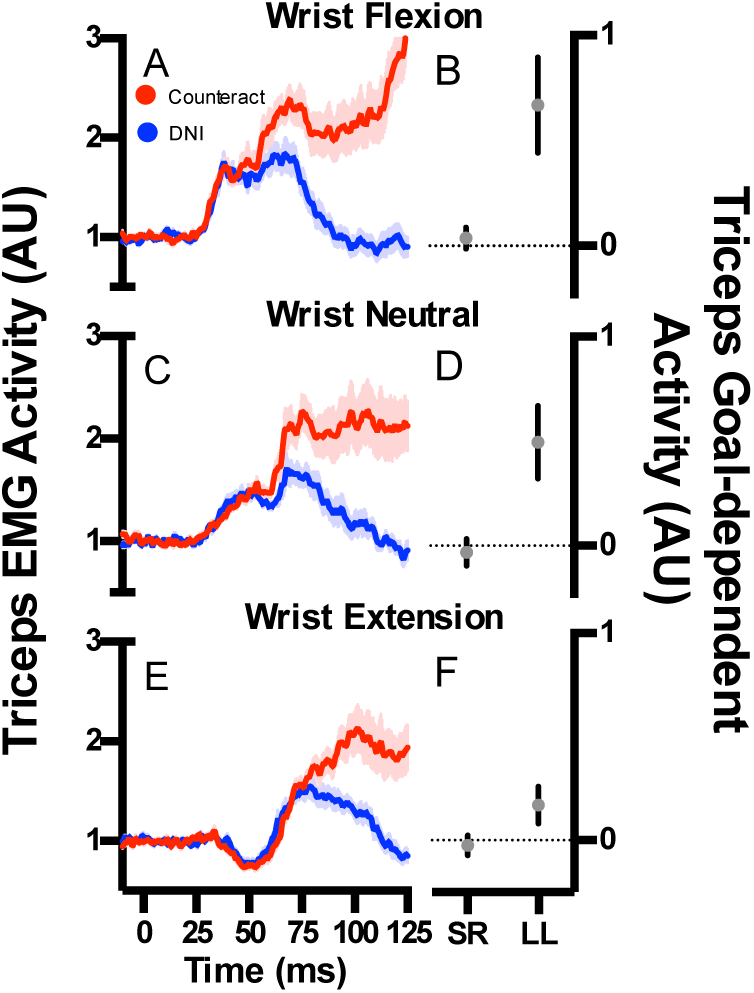
A: Mean triceps EMG activity following the mechanical perturbations that flexed the elbow and flexed the wrist. Red and blue traces reflect Counteract and Do Not Intervene blocks, respectively. Data is aligned to perturbation onset. Shading reflects ±1 SEM. B: Goal-dependent activity within the spinal (SR) and long-latency (LL) epochs for trials in which the mechanical perturbation flexed the elbow and flexed the wrist. Error bars reflect 95% confidence internals. C: Same format as A, but for trials where the elbow was flexed and no perturbation was applied to the wrist. D: Same format as B, but for trials where the elbow was flexed and no perturbation was applied to the wrist. E: Same format as A, but for trials where the mechanical perturbations flexed the elbow and extended the wrist. F: Same format as B, but for trials where the mechanical perturbations flexed the elbow and extended the wrist.

### The spinal stretch reflex accounts for arm orientation

The observation in our first experiment, that the triceps’ spinal stretch reflex was tuned to the hand’s displacement, may have reflected hardwired connections from wrist afferents to triceps motorneurons. We ruled out this possibility in a second experiment by showing that the tuning was diametrically altered when participants changed the orientation of their arm.

Participants (n = 15) completed one block of trials by grasping the robot’s handle naturally, with their thumb pointing upwards (i.e., Upright) and a second block by grasping the handle with their thumb pointing downwards (i.e., Flipped; **Fig. 4A**,**B**). For both blocks of trials, participants placed their hand on a central target. After a brief delay the robot flexed their elbow, and simultaneously either flexed or extended their wrist. Participants were instructed to counteract the perturbation by returning their hand to the target quickly and accurately. Critically, the different arm orientations dissociated how wrist rotation translated to hand movement relative to the target. As a result, wrist flexion perturbations displaced the hand further from the target when participants adopted the Upright compared to the Flipped orientation, and wrist extension perturbations displaced the hand further from the target when participants adopted the Flipped compared to the Upright orientation (**Fig. 4F**,**G**).

**Figure 4.**
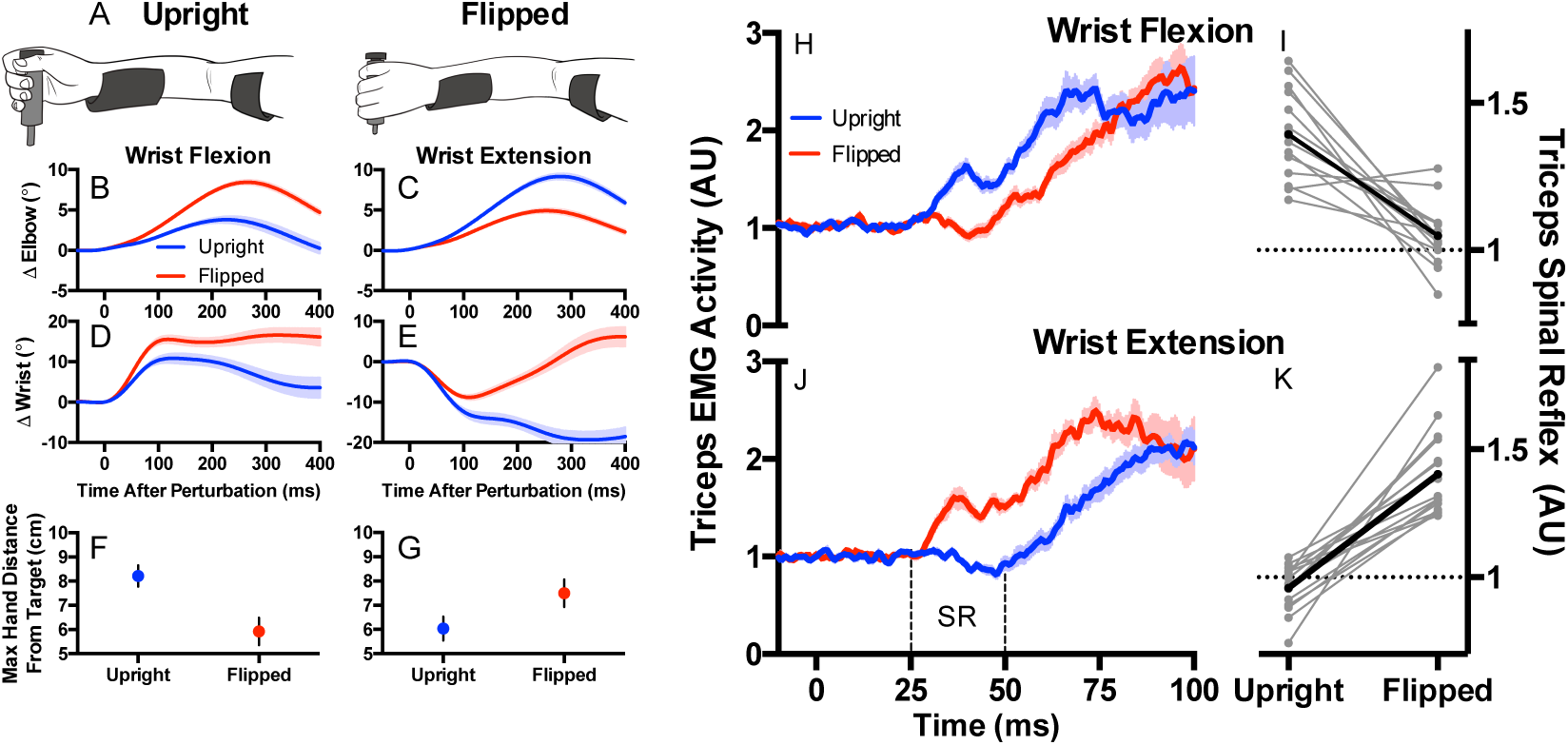
A: Cartoon of the Upright and Flipped orientations. B: Mean change in elbow angle following perturbations that flexed the elbow and flexed the wrist. Blue and red traces reflect the Upright and Flipped arm orientations, respectively. Data aligned to perturbation onset. Shading reflects ±1 SEM. C: Same format as B but for perturbations that extended the wrist. D: Same format as A but for mean change in wrist angle. E: Same format as D but for perturbations that extended the wrist. F: Maximum distance the hand was displaced from the target following perturbations that flexed the elbow and flexed the wrist. Blue and red dots reflect the group mean for the Upright and Flipped arm orientations, respectively. Error bars reflect ±1 SEM. G: Same format as F but for perturbations that extended the wrist. H: Mean triceps EMG activity for perturbations that flexed the elbow and flexed the wrist. Blue and red traces reflect the Upright and Flipped arm orientations, respectively. Data aligned to perturbation onset. Shading reflects ±1 SEM. I: Mean triceps EMG activity in the spinal stretch reflex epoch (SR: 25-50 ms relative to perturbation onset) when the wrist was flexed as a function of the Upright and Flipped Orientations. Thin grey lines reflect individual participants whereas the thick black line reflects the group mean. J: Same format as H, but for trials when the wrist was extended. K: Same format as I, but when the wrist was extended.

Participants readily changed how they coordinated their elbow and wrist joints as a function of arm orientation (**Fig. 4B-E**). Strikingly, the triceps’ spinal stretch reflex was again tuned to the hand’s displacement from the target rather than the elbow’s rotation (wrist flexion, t(14) = 6.05, p <0.001; wrist extension, t(14) = -8.66, p < 0.001). In fact, changing the arm’s orientation diametrically altered the pattern of the triceps’ spinal stretch reflex and did so in a way that was appropriate for returning the hand to its initial location (**Fig. 4H-K**).

## Discussion

The spinal stretch reflex is generated exclusively by spinal circuitry (Liddell and Sherrington, 1924) and is typically thought to re-establish the initial length of a muscle when it is unexpectedly stretched by an external disturbance (Easton 1972; Kandel et al, 2013). Regulating the length of individual muscles is the simplest way to stabilize the body against disturbances and such a control scheme could be implemented by monosynaptic and homonymous connections between muscle spindles and motorneurons that arise from and target the same muscle – the typical description of the architecture of the spinal feedback pathway (Chen et al, 2003; Kandel et al, 2013).

Our results reveal that the spinal feedback pathway that generates the spinal stretch reflex produces a more sophisticated control solution when stabilizing the hand in the presence of external disturbances. The triceps’ spinal stretch reflex does not merely reflect the local stretch of the triceps and, as such, does not act to locally regulate the length of elbow muscles. Rather, it integrates information from both the elbow and wrist (Exp. 1), and even takes into account the arm’s orientation (Exp. 2), in a manner that supports postural control of the hand – that is, maintaining the hand at its pre-perturbation location. Thus, this spinal feedback pathway can exploit the arm’s musculoskeletal redundancy and can implement sophisticated control laws (Todorov 2004; Todorov and Jordan, 2002) usually considered unique to a transcortical feedback pathway (Scott, 2004; Scott, 2016). This is not to say that this spinal feedback pathway can do everything a transcortical feedback pathway can do. Consistent with many studies, we found that the triceps’ spinal stretch reflex is not modulated by verbal instructions about how to respond to the applied perturbation (Exp. 1b). The response we document here is similar to other functional reflexes mediated entirely by spinal circuitry (e.g., withdrawal; crossed extensor; wiping reflexes), in that the ultimate motor output, although complex and purposeful, is tightly coupled to the sensory input and does not appear to have the ability to arbitrarily transform sensory feedback into any desired action.

The neural implementation of the sophisticated control laws that support postural hand control requires flexibly combining simultaneous inputs from both homonymous (i.e., triceps) and heteronymous muscles (i.e., muscles that span the wrist). The presence of heteronymous connections between arm muscles is well established, including wrist and elbow muscles as well as elbow and shoulder muscles (Cavallari and Katz 1989; Desillingly and Burke, 2012; Manning and Bawa, 2011; McClelland, Miller and Eyre, 2001). Interestingly, the fastest spinal feedback pathway appears not to always take advantage such heteronymous connections. For example, previous studies focusing primarily on whole arm reaching have specifically noted that spinal stretch reflexes at the shoulder and elbow respond only to local muscle stretch even when integrating information from the other joint would aid task performance (Kurtzer et al, 2008; Kurtzer et al., 2009; Kurtzer et al., 2014; Pruszynski et al, 2011; Soechting and Lacquiniti, 1988). Why would heteronymous connections functionally link the elbow and the wrist, but not the elbow and the shoulder? One possibility is that such differences in neural control arise because of differences in how these joints are anatomically arranged. Unlike the upper arm and forearm, the forearm and hand are usually aligned with one another meaning that, for keeping the hand stable in a circumscribed part of external space, small disturbances at the wrist can be naturally opposed by counter-rotations at the elbow.

Our findings also reveal that this spinal feedback pathway has a mechanism that tunes the inputs from heteronymous connections such that changing the arm’s orientation diametrically alters how the spinal reflex at the elbow integrates information arising from the wrist joint. This non-linear mapping between sensory inputs and motor outputs increases the computational capacity of this circuit and seems likely to be implemented via presynaptic inhibition that selectively gates which heteronymous inputs influence triceps’ motor neurons. Specifically, the presynaptic inhibition could regulate whether the stretched heteronymous wrist muscle provides the triceps’ motorneuron excitatory input through a monosynaptic direct pathway, or inhibitory input via an indirect pathway routed through an inhibitory interneuron (**Fig. 5**). Recent work in the mouse has shown that a specific set of GABAergic spinal interneurons exerts presynaptic inhibitory control of incoming afferent feedback, and ablation of these neurons has detrimental consequences for movement execution (Fink et al, 2014). We speculate that this same class of spinal interneurons underlies the selective gating of heteronymous connections critical to the sophisticated spinal feedback control we describe here.

**Figure 5.**
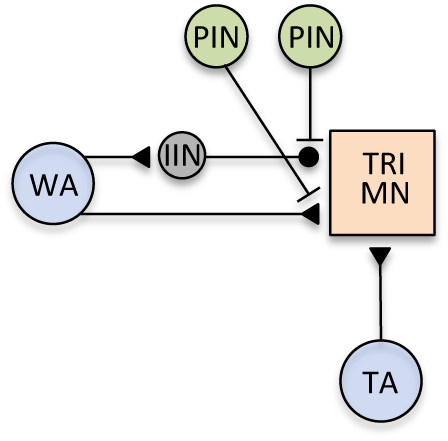
Cartoon of proposed spinal circuit. The stretched homonymous triceps muscle (TA) and the stretched heteronymous wrist muscle (WA) provide excitatory input into the triceps’ spinal motorneurons (TRI MN). Inhibitory input into the TRI MN is also provided from the stretched heteronymous wrist muscles (WA) through an inhibitory interneuron (IIN). Separate pools of presynaptic inhibitory neurons (PIN) selectively gate whether the excitatory or inhibitory input from the heteronymous wrist muscle exert their influence on the triceps’ motorneuron. This circuit architecture predicts that inhibitory stretch reflexes should show an initial brief moment of excitation, because the indirect inhibitory input from the WA is slightly delayed relative to the direct excitatory input from the TA (see initial excitation in blue trace of Fig 2A and red trace of Fig 4H).

## Methods

### Participants

Twenty-five individuals volunteered for *Experiment 1* (15 males, 10 females) and 15 individuals volunteered for *Experiment 2* (6 males, 9 females). All participants reported having normal or corrected-to-normal vision and provided informed written consent prior to data collection. This study was approved by the Office of Research Ethics at Western University and was conducted in accordance with the Declaration of Helsinki.

### Apparatus

Participants grasped the handle of a three degree-of-freedom (shoulder, elbow and wrist) exoskeleton robot. The robot allows for flexion and/or extension of the shoulder, elbow and/or wrist in a horizontal plane, and is equipped with motors that generate flexion or extension loads at these joints and encoders to measure joint kinematics. Visual stimuli were presented downward by a 46- inch LCD monitor (60 Hz, 1,920 × 1,080 pixels, Dynex DX-46L262A12, Richfield, MN) onto a semi-silvered mirror that occluded vision of the participant’s arm. Participants were comfortably seated in a height adjustable chair and the lights in the experimental suite were extinguished for the duration of data collection.

### Experimental procedures

Participants controlled a cursor (turquoise circle: 1 cm diameter) that was mapped to the position of the robot’s handle. Participants began each trial by moving the cursor to a start-position (red circle: 1 cm diameter) and maintained this position for 500 ms. The robot then gradually applied increasing loads that flexed the elbow and wrist for 1500 ms, which plateaued at 2 Nm and 1 Nm, respectively (i.e., the pre-load). A target (red circle: 1 cm diameter) was then presented approximately 5 cm in front of the start-position, which corresponded to the position of the cursor when the participant’s shoulder, elbow and wrist were at 70°, 60° and 10° of flexion, respectively (external angle coordinate system). When the participant moved the cursor to the target and had their wrist between 5 and 15° of flexion, the target changed from red to green and the start-position was extinguished. The cursor was extinguished and the target changed to green to yellow – which served as a perturbation warning cue – after participants maintained the cursor at this location with the wrist in the required configuration for 1000 ms. Following a randomized foreperiod (1000-2500 ms) the robot then applied a 2 Nm step-torque (i.e., the perturbation) at the elbow that flexed the elbow joint, and simultaneously applied a 1 Nm, -1 Nm perturbation or no perturbation at the wrist. The perturbation condition was randomized on a trial-by-trial basis.

Ten participants from *Experiment 1* were instructed to counteract the perturbations and return the cursor to the target as quickly as possible. These ten participants completed 100 trials of each of the three experimental conditions (1: an elbow flexion perturbation paired with a wrist flexion perturbation; 2: an elbow flexion perturbation paired with no wrist perturbation; 3: an elbow flexion perturbation paired with a wrist extension perturbation) in a randomized order, totaling 300 trials. The remaining 15 participants from *Experiment 1* completed 75 trials for each of three aforementioned experimental conditions, and also completed an additional block of trials in which they were instructed to “not intervene” following the single- or multi-joint perturbations. These participants completed 75 trials for each of these additional experimental conditions, thus totaling 450 trials. The ordering of the “Do Not Intervene” and the “Counteract” blocks were randomized across these participants.

The 15 participants from *Experiment 2* completed two blocks of trials in which they were instructed to return the cursor to the target as quickly as possible following multi-joint perturbations that flexed the elbow and wrist, or that flexed the elbow and extended the wrist. Critically, the blocks in *Experiment 2* differed by how the participants physically grasped the robot handle. In one block, participants grasped the handle with their thumb pointing upward (i.e., Upright), whereas in the other block, participants internally rotated their forearm and shoulder and grasped the handle with their thumb pointing downward (i.e., Flipped: see **Fig. 4A**,**B**). These different arm orientations dictated how the wrist perturbation moved the cursor relative to the target. For example, perturbations that flexed the wrist moved the cursor away from the target when participants adopted the Upright orientation, but moved the cursor towards the target when participants adopted the Flipped orientation. Participants completed 75 trials for each of the experimental conditions across both blocks, for a total of 300 trials.

All participants were given movement feedback after each trial. The target changed from yellow to green if participants returned the cursor to the target in less than 375 ms following the single or multi-joint perturbation, or changed from yellow to red otherwise. All participants completed practice trials prior to the data collection until a success rate of approximately 75% was achieved. Rest breaks were given approximately every 20 minutes during data collection or when requested.

### Muscle activity

Participants’ skin was cleaned with rubbing alcohol and a EMG surface electrode (Delsys Bagnoli-8 system with DE-2.1 sensors, Boston, MA) contacts were coated with a conductive gel. The EMG electrode was then placed on the belly of the lateral head of the triceps brachii (a monoarticular elbow extensor) and a reference electrode was placed on participants’ left clavicle. EMG signals were amplified (gain = 1000), and then digitally sampled at 2000 Hz.

### Data reduction and analysis

Angular position of the shoulder, elbow and wrist were sampled at 500 Hz. EMG data were band-pass filtered (20 – 250 Hz, 2-pass –2^nd^ order Butterworth) and full-wave rectified. The TRI muscle activity was normalized to its own mean activity 200 ms prior to perturbation onset. Joint kinematics and EMG were recorded from -200 ms to 400 ms relative to perturbation onset, and low-pass filtered (12 Hz, 2-pass 2^nd^-order Butterworth)

We compared mean activity of the triceps spinal stretch reflex and long-latency stretch response, with repeated-measures ANOVAs or paired sample t-tests. Post-hoc contrasts were completed with within-subject contrasts (i.e., trend analysis) or with paired sample t-tests. Experimental results were considered reliably different if p < 0.05.

## Acknowledgements

This work was supported by the Natural Science and Engineering Council of Canada (NSERC Discovery Grants to JAP and PLG). JW is supported by post-doctoral fellowships from NSERC and the BrainsCAN program at Western University. JAP received a salary award from the Canada Research Chairs Program.

## Supplementary Figure

**Figure 1:**
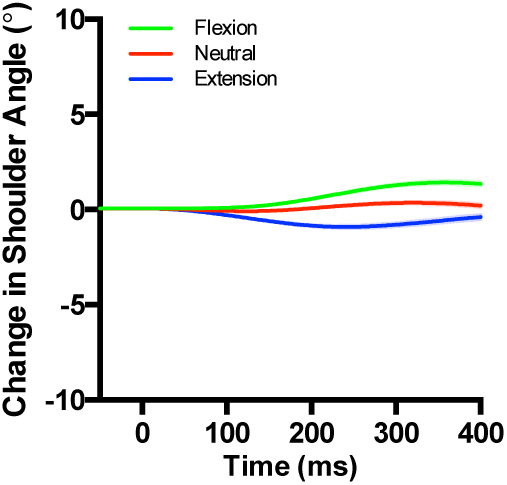
Mean change in shoulder angle aligned to perturbation onset. Green, and blue traces reflect perturbations that flexed and extended the wrist, respectively. Red trace reflects trials in which no mechanical perturbation was applied to the wrist. Shading reflects ±1 SEM.

